# Differential modulation of cellular phenotype and drug sensitivity by extracellular matrix proteins in primary and metastatic pancreatic cancer cells

**DOI:** 10.1101/2022.11.11.516201

**Authors:** Olalekan H Usman, Sampath Kumar, Reddick R Walker, Gengqiang Xie, Hyeje Sumajit, AbdelAziz R. Jalil, Subramanian Ramakrishnan, Lawrence J Dooling, Yue Julia Wang, Jerome Irianto

## Abstract

Pancreatic cancer adenocarcinoma (PDAC) is reported to be the third highest cause of cancer-related deaths in the United States. PDAC is known for its high proportion of stroma, which accounts for 90% of the tumor mass. The stroma is made up of extracellular matrix (ECM) and non-malignant cells such as inflammatory cells, cancer-associated fibroblasts, and lymphatic and blood vessels. Here, we decoupled the effects of the ECM on PDAC cell lines by culturing cells on surfaces coated with different ECM proteins. Our data show that the primary tumor-derived cell lines have different morphology depending on the ECM proteins on which they are cultured, while metastatic lesion-derived PDAC lines’ morphology does not change with respect to the different ECM proteins. Similarly, ECM proteins modulate the proliferation rate and the gemcitabine sensitivity of the primary tumor PDAC cell lines, but not the metastatic PDAC lines. Lastly, transcriptomics analysis of the primary tumor PDAC cells cultured on different ECM proteins reveals the regulation of various pathways, such as cell cycle, cell adhesion molecules, and focal adhesion, including the regulation of several integrin genes that are essential for ECM recognition.

## Introduction

Pancreatic cancer adenocarcinoma (PDAC) is reported to be the third highest cause of cancer-related deaths in the United States and it is predicted to replace colorectal cancer as the second highest before 2030 ^1,2^. Although in general, there has been a downward trend in the number of cancer deaths—a 32% decrease from peaks in 1991 and 2019 ^3^—PDAC has been an exception, with less than 5% of patients surviving for 5 years ^4^. This poor prognosis has been attributed to the difficulty of early-stage diagnosis due to the lack of specific tumor markers, along with resistance to radiotherapy, chemotherapy, and molecular targeted therapy ^5^. PDAC is known for its high proportion of stroma, which accounts for 90% of the tumor mass ^6^. The stroma is made up of extracellular matrix (ECM) and non-malignant cells such as inflammatory cells, cancer-associated fibroblasts (CAF), and lymphatic and blood vessels. These stromal components interact bidirectionally and dynamically with the tumor cells ^7^. Tumor fibrotic stroma has been a major contributor to PDAC progression and invasion, together with resistance to treatment ^8,9^. The resulting stiffness from high stroma deposition in PDAC can also result in deformation of the vasculature, thereby limiting oxygen, nutrients, and drug transport to the tumor cells ^10^.

The ECM is a highly organized three-dimensional meshwork that surrounds and provides physical support for the cells ^11^. ECM is mainly made up of glycoproteins and fibrous proteins including collagens, fibronectin, laminin, vitronectin, proteoglycans, and elastin ^12,13^. The reorganization, density, posttranslational modifications, and remodeling of its constituent proteins confer the ECM the ability to respond to both biochemical and mechanical cues from the tumor microenvironment^14^. In this study, we focus on collagen I, fibronectin, laminin, and vitronectin. Of the 28 collagens that have been identified, collagen I is known to promote tumorigenesis, and immunohistochemistry staining from different studies showed a significantly high level of the protein in both mouse and human PDAC samples ^15,16^. Collagen I and collagen III make up approximately 80% of the ECM mass ^17^. The crosslinking of collagen I and its remodeling by proteases have been implicated in tumor progression processes including invasion ^18^ and resistance to cell death ^19^. In addition, Olivares and colleagues showed that collagen can be a source of proline (one of its components), which can serve as fuel for cancer cells when other nutrients are limited ^15^. Fibronectin is a key glycoprotein in the ECM, and it has been reported that fibronectin is required for the assembly and deposition of other ECM proteins including collagen I ^20^. Fibronectin contributes to tumor progression and dormancy by increasing proliferation signaling via integrin clustering ^21^, inhibiting apoptosis ^22^, and promoting the formation of a pre-metastatic niche ^23^. In addition, in small-cell lung cancers, fibronectin has been reported to promote chemoresistance to drugs such as etoposide and doxorubicin ^23,24^. Laminin is involved in providing tensile strength in the basement membrane, which helps to maintain the apical-basal polarity of epithelial structures ^25^. Different groups have shown that laminin expression correlates with tumor invasion ^26 27^. In a similar manner to other ECM proteins mentioned above, vitronectin acts as an adhesive, connecting the cells to the matrix through integrins. A high level of vitronectin in neuroblastoma correlates with poor prognosis ^28^.

Here, we describe the results of decoupling the effects of the ECM on PDAC cell lines by culturing the cells on surfaces coated with different ECM proteins. Our data show that primary tumor-derived cell lines have different morphologies and proliferation rates that depend on the ECM proteins on which they are cultured, while metastatic lesion-derived PDAC lines’ morphology and proliferation rate do not change with respect to different ECM proteins. Next, we examined how these ECM proteins affect cell sensitivity to gemcitabine’s cytotoxicity. Similar to the cellular morphological and proliferation rate findings, ECM proteins significantly modulate the gemcitabine sensitivity of primary tumor PDAC cells, while the modulation in the metastatic lines was limited. Transcriptomics analysis shows differential expression of genes involved in pathways such as cell cycle, DNA replication, focal adhesion, MAPK signaling pathway, and more when primary tumor PDAC cells were cultured on different ECM proteins. We observed differential regulations of ECM receptor integrins between the cells cultured on fibronectin and vitronectin. Finally, we also observed the differential response to ECM proteins when the cells were cultured in 2D and 3D cultures.

## Results

### ECM proteins have different effects on PDAC cell lines’ morphology and proliferation

To study the effect of ECM proteins on the morphology of cells, primary tumor PDAC cell lines (MIA PaCa-2 and PANC-1, **Figure S1A**) were seeded at approximately 3200 cells/cm^2^ and cultured for 6 days in well plates coated with collagen I, fibronectin, laminin, and vitronectin using culture media supplemented with 2% B27. We supplemented the culture media with serum-free supplement B27 instead of the conventional fetal bovine serum (FBS) mainly because FBS consists of albumin, growth factors, and other proteins including the ECM proteins, like fibronectin. These can be deposited on the culture surface and interact with the cells during culture, which risked confounding our observations ^29^. Microscopy imaging shows that MIA PaCa-2 spreads in wells coated with fibronectin and vitronectin similarly to cells grown in 10% FBS media. Likewise, MIA PaCa-2 cells are rounded in collagen I and laminin in a manner similar to cells grown on a plate with no ECM protein coating (**Figure 1Ai top**). The PANC-1 line adheres and spreads on collagen I, fibronectin, and vitronectin substrates, and they were rounded on laminin **(Figure 1Ai bottom)**. We quantified the morphology of the cells grown on each ECM protein by measuring the aspect ratio of the cells. Our quantification supports the observations above: The aspect ratio is higher for MIA PaCa-2 cells cultured on fibronectin and vitronectin **(Figure 1Aii)** and for PANC-1 cells cultured on collagen I, fibronectin, and vitronectin **(Figure 1Aiii)**.

**Figure 1.**
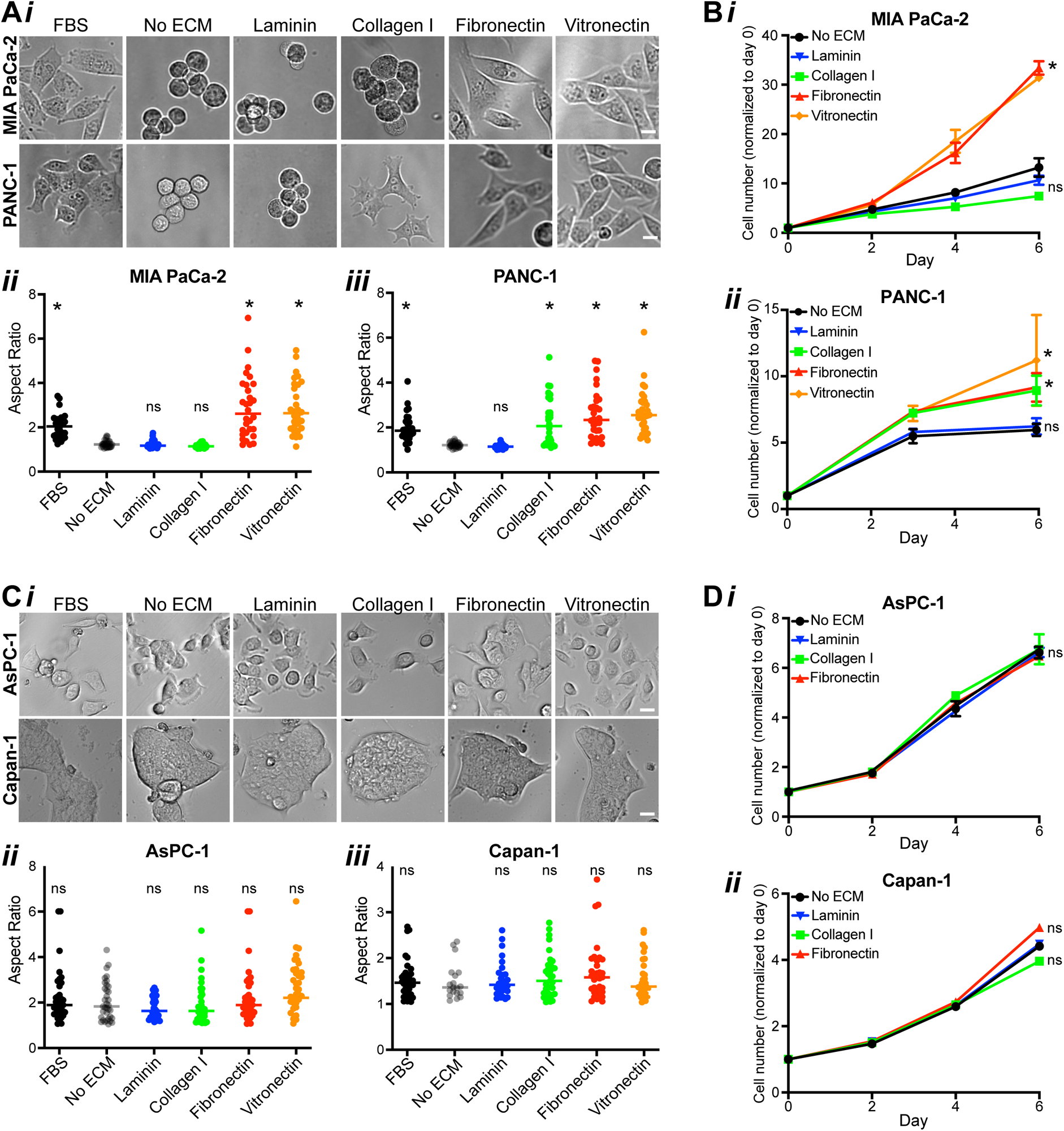
ECM proteins have different effects on pancreatic ductal adenocarcinoma (PDAC) cell lines morphology and proliferation. **(A)** The primary tumor-derived cell lines, MIA PaCa-2 and PANC-1, were cultured in media supplemented with either FBS or the serum-free supplement 2% B27. To investigate the effect of ECM proteins, the cells cultured in 2% B27 were seeded on surfaces without (no ECM) or with ECM protein coating: laminin, collagen I, fibronectin, and vitronectin. Representative images show that MIA PaCa-2 cells (top) adhere and spread in FBS and on surfaces coated with fibronectin or vitronectin, while they have rounded morphology on no ECM, laminin, and collagen I surfaces **(i)**. The cellular aspect ratio (AR) is significantly higher in MIA PaCa-2 cells cultured on fibronectin and vitronectin **(ii)**. PANC-1 cells (bottom) also spread in FBS and on surfaces coated with collagen I, fibronectin, and vitronectin, while they remain rounded in no ECM and laminin-coated surfaces. The AR for PANC-1 is significantly higher on surfaces coated with collagen I, fibronectin, and vitronectin **(iii)** (N = 3 experiments, n = 30 cells per group, \**p* < 0.01 vs. no ECM, ns: *p* > 0.05 vs. no ECM, scale bar: 15µm). **(B)** MIA PaCa-2 and PANC-1cell lines were cultured on surfaces coated with collagen I, fibronectin, laminin, and vitronectin. The number of cells under each condition was quantified on days 2, 4, and 6 using a luminescent cell viability assay. The number of cells on days 2, 4, and 6 was normalized to the number of cells seeded. MIA PaCa-2 has a higher proliferation rate on fibronectin and vitronectin substrates compared to collagen I, laminin, or well with no ECM coating **(i)**. PANC-1 cultured on collagen I, fibronectin, and vitronectin show a higher proliferation rate compared to those grown on laminin and no ECM **(ii)**. (N = 3 experiments, mean ± SEM, \**p* < 0.01 vs. no ECM, ns: *p* > 0.05 vs. no ECM). **(C)** Representative images of the metastatic lesion-derived cells, AsPC-1 (top) and Capan-1 (bottom), shows that their morphology do not change regardless of the ECM proteins on which they are cultured **(i)**. The cellular aspect ratio of Capan-1 and AsPC-1 is not significantly different in ECM proteins compared to no ECM **(ii-iii)** (N = 3 experiments, n = 30 cells per group, *p < 0.01 vs. no ECM, ns: *p* > 0.05 vs. no ECM, scale bar: 15µm). **(D)** The number of AsPC-1 and Capan-1 cells cultured on the different ECM coatings were quantified on days 2, 4, and 6, and normalized to the number of cells seeded. For both AsPC-1 and Capan-1, the proliferation rate of the cells cultured on the different ECM proteins was not significantly different to those cultured on no ECM plates **(i-ii)** (N = 3 experiments, mean ± SEM, ns: *p* > 0.05 vs. no ECM).

To note, we adopted the ECM coating concentrations that were commonly used in other studies and recommended by the manufacturers. However, multiple studies have reported that the degree of cell spreading is a function of ECM coating concentration^30,31^ and the response is biphasic^32–35^, where the degree of cell spreading will drop when the ECM concentration is too high. To check whether the ECM coating concentrations that we chose are appropriate for this study or not, we cultured the MIA PaCa-2 cells on surfaces coated with a range of ECM protein concentrations. For the laminin and collagen I groups, the MIA PaCa-2 cells remain rounded with the coating concentrations that are 10-fold lower and higher than the chosen concentrations **(Figure S1Bi-ii)**. For the fibronectin and vitronectin groups, the aspect ratio of MIA PaCa-2 cells decreased significantly when they were cultured on the lower coating concentration, but the aspect ratio did not increase further on the higher coating concentrations **(Figure S1Biii-iv)**. These results suggest that the coating concentrations that we chose are appropriate for our study. Another thing to note is that the degree of cell spreading observed here is a result of an extended period of culture for 6 days, which provides the cells ample time to fully adhere to the culture substrate. To investigate the effect of ECM proteins on MIA PaCa-2 adhesion capability at a shorter time scale, we performed an attachment assay, where the cells were only allowed 6 hours to adhere to the different ECM-coated surfaces. Similar to the morphology data, the number of attached cells in the laminin and collagen I groups is significantly lower when compared to the cells cultured in FBS media. While the cell count of the fibronectin and vitronectin groups are comparable to the FBS group **(Figure S1Ci-ii)**.

Next, we studied the effect of ECM proteins on the growth of the cell lines using a luminescence assay. The number of cells under each condition was quantified over days of culture using a luminescent cell viability assay and normalized to the number of cells seeded. Resonating with the morphology findings, MIA PaCa-2 cells on fibronectin- and vitronectin-coated surfaces have a significantly higher proliferation rate when compared to cells on collagen I and laminin **(Figure 1Bi)**, suggesting the correlation between cell spreading and proliferation rate. The same is the case for the spreading PANC-1 cells cultured on collagen I, fibronectin, and vitronectin: They double faster relative to those cultured on laminin **(Figure 1Bii)**.

As a comparison, we also performed the same experiments on a couple of metastatic lesion-derived PDAC cell lines, AsPC-1 and Capan-1, which were derived from the ascites^36^ and the liver^37^, respectively. Microscopy imaging reveals that these cell lines moderately spread on all ECM coatings and wells with no ECM coating **(Figure 1Ci)**. The aspect ratio quantification shows that the morphology of the cell lines does not change with respect to the different ECM proteins **(Figure 1Cii–iii)**. Capan-1 cells are often clustered together, and to quantify their aspect ratio, we stained the cells’ nuclei and filamentous actin (F-actin), which allowed us to differentiate individual cells within the clusters **(Figure S1A)**. The attachment assay for the Capan-1 also did not show any significant differences between the different ECM proteins **(Figure S1Ciii-iv)**. From the luminescence-based proliferation assay, both AsPC-1 and Capan-1 proliferation rates were not significantly influenced by any of the ECM proteins **(Figure 1Di–ii)**, even with a longer culture period **(Figure S1D)**.

### ECM proteins alter cell response to gemcitabine

Gemcitabine is an analog of deoxycytidine that inhibits the elongation step during DNA synthesis and consequently, prevents cell growth ^38,39^. It is used as the first-line treatment for PDAC patients^40^. However, after a few weeks of treatment, most patients become resistant to the drug ^41^. To assess the cytotoxic effect of gemcitabine on the cell lines **(Figure 2A)**, we treated them with gemcitabine for 72 hours and the cell numbers were quantified using a luminescence cell viability assay. Curve fitting^42^ was performed on the luminescence data to derive the gemcitabine dose-response curves and their metrics, including the half maximal inhibitory concentration (IC50) and the least percentage of surviving cells (Emax), as the indication of the drug potency and efficacy, respectively. We found that MIA PaCa-2 and Capan-1 are more sensitive to the drug treatment, where both their IC50 and Emax are significantly lower than PANC-1 and AsPC-1 **(Figure 2B).**

**Figure 2.**
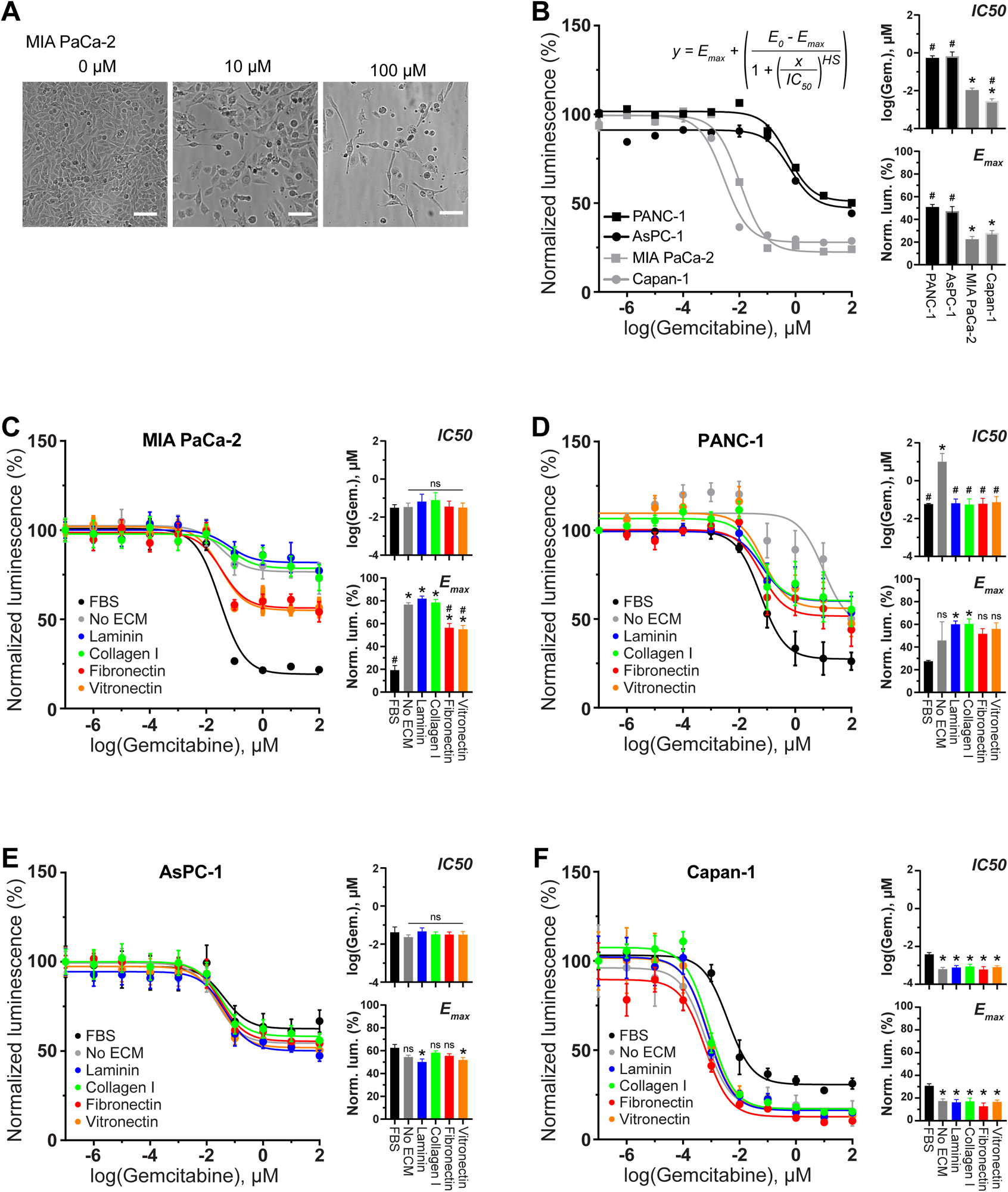
ECM proteins alter gemcitabine sensitivity of the primary-derived PDAC cell lines, but not the metastatic lines. **(A)** Cytotoxic effect of gemcitabine on MIA PaCa-2 cells was assessed by treating the cells with different concentrations of the drug. Representative images show that the number of surviving cells decreases with increasing concentrations of gemcitabine (scale bar: 100µm). **(B)** MIA PaCa-2, PANC-1, AsPC-1, and Capan-1 were treated with gemcitabine for 72 hours and the cell numbers were measured using luminescence cell viability assay. Curve fitting (inset) was performed on the luminescence data to derive the dose-response curves and the corresponding IC50 and Emax (right). The IC50 and Emax of MIA PaCa-2 and Capan-1 are significantly lower than PANC-1 and AsPC-1, suggesting that both MIA PaCa-2 and Capan-1 cells are more sensitive to the drug treatment. (N =. 3 experiments, n = 2–3 wells, \**p* < 0.05 vs. PANC-1, ^#^*p* < 0.05 vs. MIA PaCa-2). **(C)** MIA PaCa-2 cells were cultured on surfaces coated with different ECM proteins and then treated with gemcitabine to study how the ECM proteins affect drug response. No significant differences were observed from the IC50 data, however, the Emax data showed that the total number of surviving cells after gemcitabine treatment is the lowest for cells cultured in FBS media, followed by the cells cultured on fibronectin and vitronectin and the cells cultured on collagen I and laminin (N =. 3 experiments, n = 2–3 wells, \**p* < 0.05 vs. FBS, ^#^*p* < 0.05 vs. no ECM, ns: *p* > 0.05 vs. FBS). **(D)** PANC-1 cells cultured on the different ECM proteins were treated with gemcitabine. Only the IC50 of the no ECM group is significantly higher than the cells cultured in FBS. While the Emax values of all the groups are generally higher than the cells cultured in FBS media, only the cells cultured on laminin and collagen I surfaces are statistically higher (N =. 3 experiments, n = 2–3 wells, \**p* < 0.05 vs. FBS, ^#^*p* < 0.05 vs. no ECM, ns: *p* > 0.05 vs. FBS). **(E)** AsPC-1 cells cultured on the different ECM proteins were treated with gemcitabine. No significant differences were observed from the IC50 data, however, the Emax values of the cells cultured on laminin and vitronectin are statistically lower than the cells cultured in FBS media (N =. 3 experiments, n = 2–3 wells, \**p* < 0.05 vs. FBS, ns: *p* > 0.05 vs. FBS). **(F)** Capan-1 cells cultured on the different ECM proteins were treated with gemcitabine. The IC50 and Emax of the cells cultured in FBS media are significantly higher than the other groups, however, no significant differences were observed between the different ECM proteins (N =. 3 experiments, n = 2–3 wells, \**p* < 0.05 vs. FBS).

PDAC is known for its high stroma, which has been implicated in chemoresistance ^43^. Next, we tested the effect of ECM proteins on the cytotoxic effect of gemcitabine on all of the cell lines. Toward this end, we cultured the cells on surfaces coated with different ECM proteins, followed by gemcitabine treatment and the luminescence cell viability assay. For the MIA PaCa-2 cells, no significant differences were observed from the IC50 data, however, the Emax data showed that the number of surviving cells after gemcitabine treatment is the lowest for cells cultured in FBS media, followed by the cells cultured on fibronectin and vitronectin (ECM proteins in which the cell have spread morphology), and then the cells cultured on laminin, collagen I, and no ECM **(Figure 2C)**. This data suggests that the modulation of gemcitabine efficacy by the ECM proteins correlates with the cell morphology and growth response **(Figure 1A-B)**. Gemcitabine sensitivity modulation by the ECM proteins was also observed in the PANC-1 cells. The absence of ECM proteins (no ECM) significantly increased the IC50 concentration, lowering the drug potency, while the Emax values of all groups are generally higher than the cells cultured in FBS media, although statistical significance was only observed in the collagen and laminin I groups **(Figure 2D)**. However, unlike the MIA PaCa-2, there is a discrepancy in PANC-1 morphology and growth response to the gemcitabine sensitivity modulation, where the rounded and slow-growing PANC-1 cultured on laminin **(Figure 1A-B)** seems to have similar gemcitabine sensitivity than the other ECM protein groups.

In contrast to the primary tumor-derived lines, limited modulations of gemcitabine sensitivity were observed in the metastatic lesion-derived lines. For the AsPC-1 cells, no significant differences were observed in the IC50 data, and limited differences can be observed in the E_max_ values, but statistical significance was observed in the laminin and vitronectin groups when compared to the FBS group **(Figure 2E)**. For the Capan-1 cells, although the differences are limited, both the IC50 and Emax values of the FBS group are statistically higher than the other groups **(Figure 2F)**, which might be driven by differences between FBS and B27 supplemented media.

### ECM proteins result in differential gene expression

To investigate the effects of different ECM proteins on MIA PaCa-2 gene expression, RNA-seq analyses were performed on MIA PaCa-2 cells that were cultured on different ECM-coated surfaces for 6 days. MIA PaCa-2 was chosen for this initial transcriptomic analysis because it is one of the lines that showed differences in morphology, attachment, growth, and gemcitabine sensitivity with respect to different ECM proteins. The gene count across the conditions was normalized using the variance stabilizing transformation from DESeq2 ^44^. The sample clustering was visualized by plotting the first two components resulting from PCA analysis on the x and y axes respectively **(Figure 3A)**. PCA analysis showed that cells cultured on collagen I and laminin, which result in rounded morphology, cluster together with cells grown without ECM. Interestingly, cells cultured on fibronectin and vitronectin, which result in spread morphology, cluster separately from each other, including from the cells cultured in FBS. These results suggest that the ECM proteins on which cells are cultured can modulate gene expression of the cells. Next, to show the differences in the gene expression pattern of cells cultured on different ECMs, the differentially regulated genes were visualized in a heatmap **(Figure 3B)**. The sample hierarchical clustering analysis reveals that cells cultured on collagen I, laminin, and no ECM protein have a similar expression pattern, while those cultured on fibronectin, vitronectin, and FBS show differential gene expression patterns, confirming our findings from the PCA analysis. The K-mean clustering of the genes reveals four distinct clusters of genes: Cluster 1 comprises genes that are highly upregulated in cells cultured in FBS, while also generally upregulated in cells cultured on fibronectin and vitronectin. Cluster 2 comprises genes that are downregulated in cells cultured in FBS. Clusters 3 and 4 comprise genes that are upregulated and downregulated, respectively, in cells cultured on vitronectin. To investigate the regulated pathways, we performed functional annotation analysis on the genes within each cluster by using Enrichr ^45^. Cluster 1 contains genes that are upregulated in FBS, fibronectin, and vitronectin, these genes are enriched for cell cycle, DNA replication, RNA transport, spliceosome, and other pathways **(Figure 3Ci)**, consistent with the higher proliferation rate of MIA PaCa-2 cells cultured on surfaces coated with fibronectin and vitronectin **(Figure 1Bi)**. Cluster 2 consists of genes that are uniquely downregulated in FBS and are enriched in pathways involving lysosome and glycosaminoglycan **(Figure 3Cii)**. Cluster 3 is made up of genes that are uniquely upregulated in vitronectin and enriched in pathways that include MAPK signaling, cell adhesion molecules, the Wnt signaling pathway, and others **(Figure 3Ciii)**. Cluster 4 contains genes that are uniquely downregulated in vitronectin and are enriched in pathways such as terpenoid backbone biosynthesis, endocytosis, and focal adhesion **(Figure 3Civ)**.

**Figure 3.**
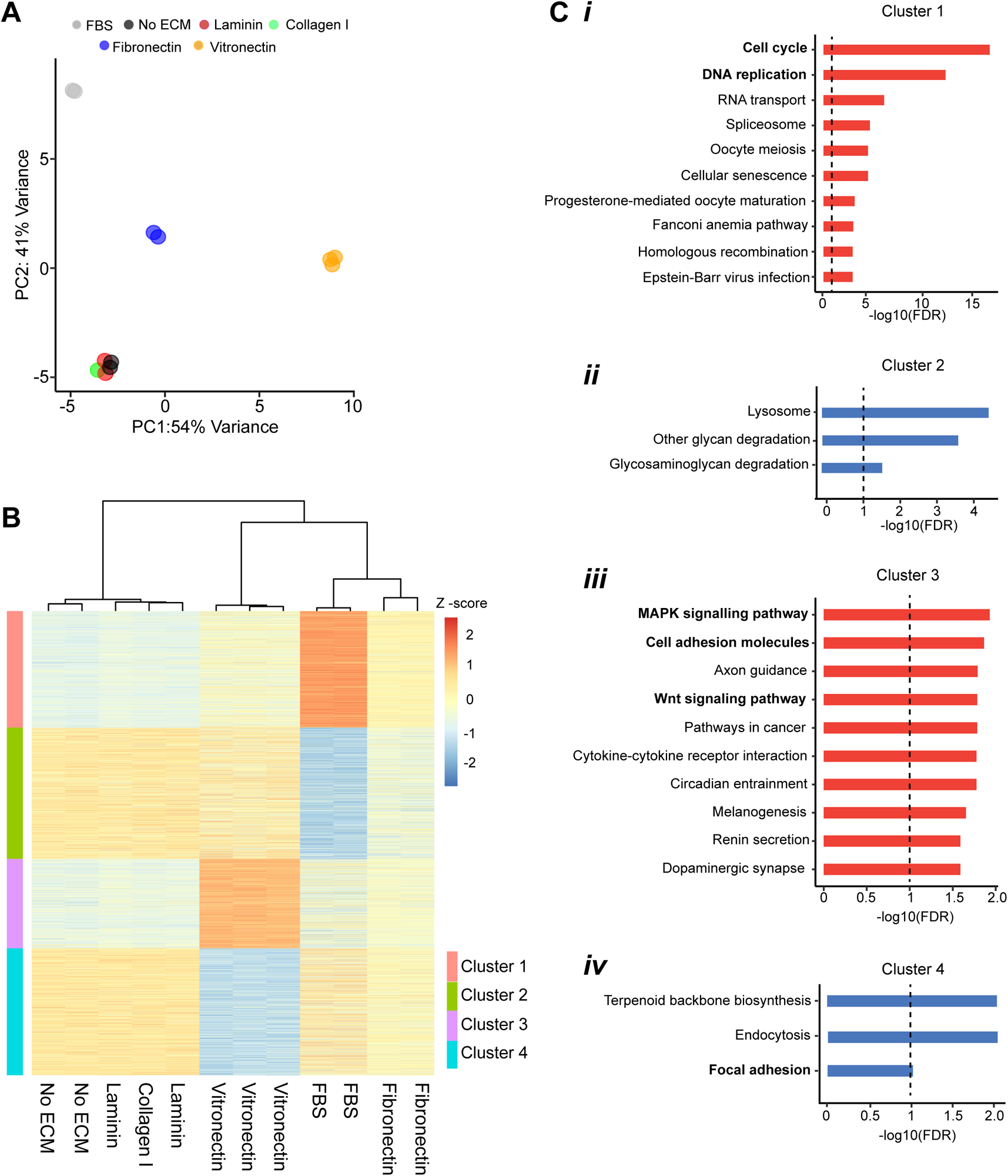
Transcriptomics analysis showing differential gene expression of MIA PaCa-2 cells cultured on surfaces coated with ECM proteins. **(A)** MIA PaCa-2 cells were cultured on different ECM proteins and subjected to RNA-seq. The gene count across the conditions was normalized using the variance stabilizing transformation from DESeq2. The sample clustering was visualized by plotting the first two components resulting from PCA analysis on the x and y axes respectively. PCA analysis showed that cells on collagen I and laminin, which result in rounded morphology, cluster together like cells grown on no ECM. Interestingly, cells cultured on fibronectin and vitronectin, which result in spread morphology, cluster separately from each other like the cells cultured in FBS. **(B)** Heatmap showing normalized read counts of the differentially regulated genes, which were derived from the comparison between MIA PaCa-2 cells cultured on the different ECM proteins and the cells cultured on the surface without ECM coating (no ECM). Clustering analysis of the heatmap reveals that cells cultured on collagen I, laminin, and no ECM protein have a similar expression pattern while those cultured on fibronectin, vitronectin, and FBS show differential gene expression patterns. Cluster 1 comprises genes that are highly upregulated in cells cultured in FBS, while also generally upregulated in cells cultured on fibronectin and vitronectin. Cluster 2 comprises genes that are downregulated in cells cultured in FBS. Clusters 3 and 4 comprise genes that are upregulated and downregulated, respectively, in cells cultured on vitronectin. Significant differentially expressed genes (adjusted p-value < 0.05, abs(log_2_FC) > 0.58) in all of the ECM proteins were combined and clustered by K-mean (with center = 4), resulting in the color-coded clusters. Hierarchical clustering was used to cluster the column using Pearson’s correlation and Ward.D2. **(C)** Bar plots showing the KEGG pathways that are enriched by the regulated genes within the clusters derived from the heatmap in Fig. 3B. The genes within each cluster were subjected to pathway analysis using Enrichr and the top 10 significant pathways were selected **(i-iv)** (FDR < 0.1). The red and blue bars indicate upregulated and downregulated pathways respectively. The dashed line on the bar plots indicates the FDR cut-off.

To investigate whether ECM proteins also regulate gene expression of the other cell lines, RNA-seq analyses were performed on PANC-1, AsPC-1, and Capan-1 cultured on fibronectin, vitronectin, and surface without ECM coating (no ECM). When compared to the no ECM group, a significant amount of genes are differentially regulated in both of the primary tumor-derived cell lines, MIA PaCa-2 and PANC-1, cultured on fibronectin and vitronectin **(Figure S2)**. However, only a very limited number of genes were differentially regulated in the metastatic lesion-derived cells, AsPC-1 and Capan-1. This observation resonates with the earlier results showing that the morphology, attachment, growth, and gemcitabine sensitivity modulations by the ECM proteins are only observed in the primary tumor-derived PDAC lines, and not in the metastatic cell lines.

Next, to reconfirm our gemcitabine sensitivity assay findings, the transcriptomic data was used to predict the gemcitabine sensitivity of MIA PaCa-2 cells cultured on the different ECM proteins, as described previously^46^. Briefly, the normalized read counts for the genes that have been shown to be positively and negatively correlated with gemcitabine sensitivity were extracted from the transcriptomic data, and then the averages of their expression from each group were used as the predictor of the sensitivity level (more details in the “Experimental procedures” section). Indeed, like the experimental findings above **(Figure 2C)**, the transcriptomic-based gemcitabine sensitivity analysis predicted that the FBS group is the most sensitive, followed by fibronectin, vitronectin, and the other groups with rounded morphology: no ECM, laminin, and collagen I **(Figure S3)**.

One of the major receptors of ECM proteins is the integrins, which are heterodimeric proteins found in the plasma membrane that connect the cells with the ECM by binding to specific recognition sites on the ECM ligands ^47,48^. In addition to focal adhesion formation, integrins are also involved in cell migration ^49^ and bidirectional biochemical and mechanical signal transduction between the cell and the ECM ^50^. MIA PaCa-2 cells express a range of integrin genes **(Figure S4)**, which potentially enable the interactions with the RGD motif within fibronectin and vitronectin through α5β1, αVβ1, αVβ5, αVβ6, and αVβ8 integrins; with the laminin through α3β1, α6β1, α7β1, α10β1, and α6β4 integrins; and with the collagen I through α10β1 integrin ^47,48^.

Both the PCA analysis **(Figure 3A)** and the clustering analysis of the differentially expressed genes **(Figure 3B)** show that MIA PaCa-2 cells respond differently when cultured on fibronectin and vitronectin. This prompted us to take a closer look at the genes that are co-regulated and uniquely regulated between these two ECM proteins. For the upregulated genes, 167 were upregulated in both fibronectin and vitronectin cultured cells, while 868 and 126 were uniquely upregulated in vitronectin and fibronectin, respectively **(Figure 4Ai)**. For the downregulated genes, 335 were downregulated in both fibronectin and vitronectin, while 1265 and 297 were uniquely downregulated in vitronectin and fibronectin, respectively **(Figure 4Aii)**. Distinct signaling pathways were enriched by each of the gene lists above **(Figure S5A-C)**. The different gene expression regulation by fibronectin and vitronectin was also observed in PANC-1 **(Figure S5D)**. Interestingly, we observed differential expression of several integrin genes in MIA PaCa-2 cells cultured on fibronectin and vitronectin (**Figure 4B)**. For example, both *ITGB8* and *ITGA6* were downregulated in cells cultured on fibronectin and vitronectin. *ITGA10* was uniquely downregulated in the fibronectin group, while *ITGB4* and *ITGB6* were uniquely upregulated in cells cultured on the vitronectin surface. This suggests that ECM proteins can regulate integrin expression levels.

**Figure 4.**
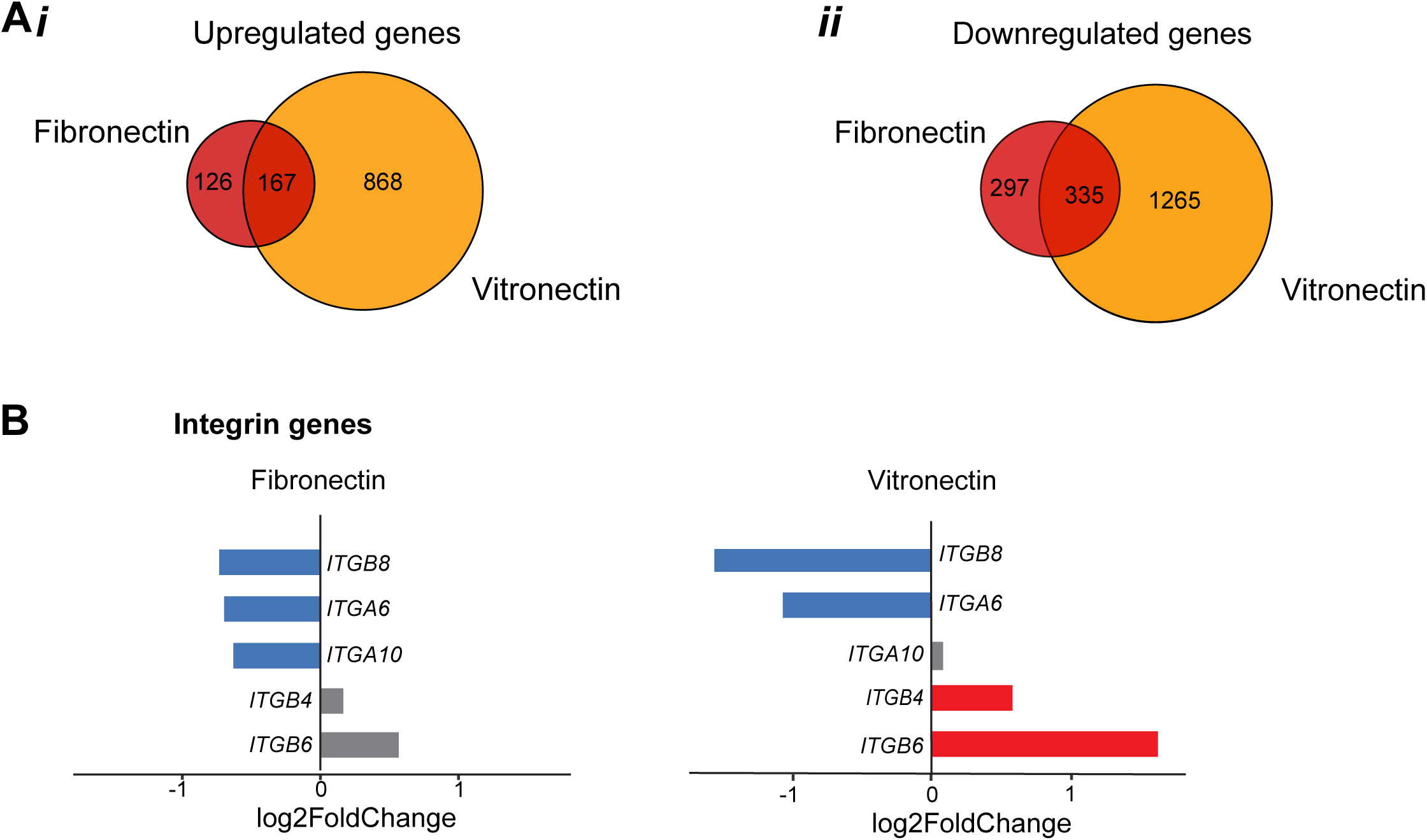
MIA PaCa-2 cells respond differently to fibronectin and vitronectin at the gene expression level. **(A)** Venn diagram showing the number of differentially expressed genes (vs. no ECM) that are regulated in both fibronectin and vitronectin (overlap), and also the genes that are unique to fibronectin or vitronectin. The left panel **(i)** shows that 167 genes are upregulated in cells cultured on fibronectin and vitronectin, while 868 and 126 genes are uniquely upregulated in vitronectin and fibronectin respectively. The right panel **(ii)** shows that 335 genes are downregulated in cells grown on fibronectin and vitronectin, while 1265 and 297 genes are downregulated uniquely in vitronectin and fibronectin respectively. **(B)** Bar plots showing the fold change of integrins in MIA PaCa-2 cell line cultured in fibronectin and vitronectin. Red, blue, and grey bars indicate upregulated, downregulated, and non-significant genes respectively.

### ECM proteins do not modulate MIA PaCa-2 morphology and growth in 3D culture

It is important to note that the findings above are limited to 2D culture, and the cellular responses to the ECM proteins might be different for cells cultured in a 3D scaffold. To facilitate the 3D culture of these cells, we adopted the polyethylene glycol (PEG) hydrogel culture^51^. We chose PEG as it is an inert material and the fabrication method enables covalent incorporation of specific ECM proteins into the gels. First, we tested the functionality of our PEG gels by culturing patient-derived PDAC organoids **(Figure 5A)**. The organoids in the PEG gels have similar morphology and grow at a similar rate to the organoids cultured in Matrigel, which is the commonly used material for PDAC organoid culture. This suggests that our PEG gels are functional and can be used to culture these organoids. Next, to confirm the covalent incorporation of ECM proteins into the PEG gels, rhodamine-tagged fibronectin or laminin was introduced to the PEG gels during the fabrication process. The rhodamine signals were clearly observed inside the PEG gels **(Figure 5B)**, confirming the incorporation of the rhodamine-tagged ECM proteins. Furthermore, to test the functionality of the incorporated ECM proteins, we seeded patient-derived glioblastoma (GBM) neurospheres into the PEG gels with and without laminin incorporation. These neurospheres are typically cultured in suspension for growth and when they are cultured in Matrigel, the cells will readily invade into the surrounding Matrigel, which is mostly made of laminin. Hence, if the incorporated laminin in the PEG gels is functional, the GBM cells should also invade the surrounding PEG-laminin gels, like in the Matrigel. Indeed, the GBM neurospheres grow and stay in place within the PEG gels without laminin, and within the PEG-laminin gels, we can clearly observe the invasion of the GBM cells into their surroundings **(Figure 5C)**, confirming the functionality of the laminin incorporated into the PEG gels. We then introduce MIA PaCa-2 cells into PEG gels with the incorporation of the different ECM proteins, followed by culture for multiple days. Unlike the 2D counterpart, we did not observe morphological **(Figure 5D)** and proliferation rate **(Figure 5E)** differences between the different ECM protein groups. This suggests that MIA PaCa-2 cells respond to ECM proteins differently when they are cultured in 2D and 3D.

**Figure 5.**
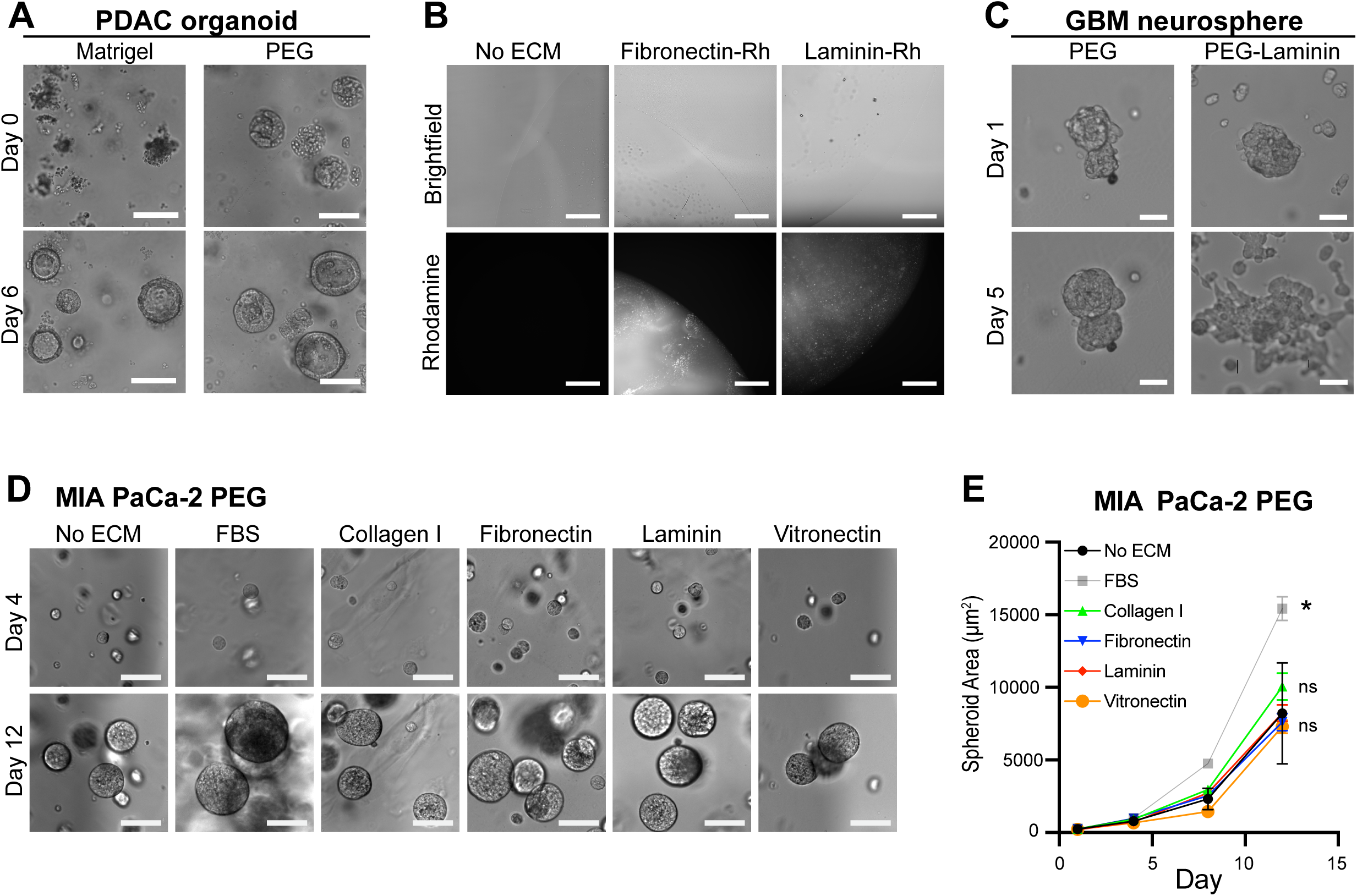
MIA PaCa-2 cells grow at a similar rate within PEG hydrogel containing various ECM proteins. **(A)** Patient-derived PDAC organoid PDM39 was cultured in Matrigel and PEG hydrogel for 6 days. Representative images show that the PDAC organoids can grow within both Matrigel and PEG hydrogel (scale bar: 100µm). **(B)** Representative fluorescence images showing the incorporation of rhodamine-tagged ECM proteins into the PEG hydrogel (scale bar: 250µm). **(C)** Patient-derived glioblastoma organoid PDM121 was cultured in PEG hydrogel without ECM incorporation (PEG) and PEG hydrogel with laminin (PEG-Laminin). Representative images showing the invasion of PDM121 only in the PEG-Laminin hydrogel, suggesting the functionality of the laminin incorporated into PEG-Laminin (scale bar: 50µm). **(D)** Representative images showing the growth of MIA PaCa-2 spheroids within PEG gels conjugated with different ECM proteins. Differences in spheroid morphology and cellular invasion were not observed between the different PEG-ECM hydrogels (scale bar: 100µm). **(E)** MIA PaCa-2 cells embedded in PEG gels conjugated with ECM proteins were cultured for 12 days and the growth was quantified by measuring the cross-section area of the spheroids. The plot shows that the MIA PaCa-2 spheroids grow faster when cultured in FBS; however, the spheroids cultured within the PEG gels with various ECM proteins grow at a similar rate as the spheroids embedded in the PEG gels without any ECM proteins (no ECM) ((N =. 3 experiments, n = 30 spheroids per group, \**p* < 0.05 vs. no ECM, ns: *p* > 0.05 vs. no ECM).

## Discussion

Phenotypic variations between different cell lines of a given cancer are common knowledge and have been reported in multiple studies ^52–54^, creating the need for investigators to choose cancer cell lines appropriate for their research questions. Our findings suggest that this is also the case for PDAC cell lines: We show that cell lines with various backgrounds respond differently to ECM proteins. In brief, significant differences in cell spreading, attachment, proliferation, gemcitabine sensitivity, and transcriptomic profile were observed when the primary tumor-derived lines, MIA PaCa-2 and PANC-1, were exposed to the different ECM coatings. Meanwhile, the differences in the metastatic lesion-derived lines, AsPC-1 and Capan-1, were very limited in comparison. There are many possible mechanisms behind these differential responses. For example, the primary tumor cell lines might express a limited set of ECM receptors --- the integrins, that only allow for adhesions to specific ECM proteins. Alternatively, it is also possible that the metastatic cell lines (AsPC-1 and Capan-1) actively secrete ECM proteins themselves, enabling them to adhere and spread on surfaces without ECM coatings.

MIA PaCa-2 cells spread and proliferate more on surfaces coated with fibronectin and vitronectin, but not on collagen I and laminin. Interestingly, the gemcitabine sensitivity and the transcriptomics also differ in different ECMs **(Figures 2C and 3)**. From the gemcitabine sensitivity assay, we observed that the spreading MIA PaCa-2 cells on fibronectin and vitronectin are more sensitive to gemcitabine than the rounded cells in the no ECM, collagen I, and laminin groups. Gemcitabine is a nucleoside analog that disturbs and inhibits DNA synthesis; considering the drug’s mode of action, it is plausible that the cells cultured on fibronectin and vitronectin are more susceptible to the drug simply because of their higher proliferation rate, i.e., higher DNA synthesis activities. However, it is also possible that the presence of ECM proteins regulates other biological processes that modulate the cells’ response to gemcitabine ^55^. From the transcriptomic study, we observed that the transcriptomic profiles of MIA PaCa-2 cultured on fibronectin and vitronectin were distinct from the no ECM, collagen I, and laminin groups. The cell cycle and DNA replication pathways were both enriched in the upregulated genes in FBS, fibronectin, and vitronectin, resonating with the increased cell proliferation rate observed in Figure 1. Additionally, the transcriptomic-based gemcitabine sensitivity predictions also agree with the pattern of MIA PaCa-2 gemcitabine sensitivity assay results. All combined, these results suggest functional transcriptomic regulation by the ECM proteins that resulted in significant changes in cellular phenotypes and drug response.

Moreover, considering the importance of the integrins as ECM receptors, MIA PaCa-2 cells express a subset of integrins **(Figure S4)**, which may partially explain their inability to adhere to collagen I, as they express only one possible integrin heterodimer, α10β1, that can recognize collagen I. The inability of MIA PaCa-2 to spread on the laminin surface is intriguing, considering the abundant expression of *ITGA3*, *ITGA6*, and *ITGB4* that can potentially bind to laminin. This discrepancy might be explained by the complex activation and inhibition of integrins by multiple proteins ^56,57^ or by the competitive pairings of the alpha and beta subunits that preferentially favor the assembly of a particular set of integrin heterodimers ^58^. Interestingly, even though fibronectin and vitronectin are recognized by a similar set of integrins that can bind to the RGD motifs ^47,48^, we observed differences in the genes regulated by fibronectin and vitronectin. Moreover, several integrin genes were also differently regulated between the cells cultured on fibronectin and vitronectin **(Figure 4B)**. The activation of integrins is known to regulate many cellular processes^47^. For example, multiple integrins have been shown to regulate cellular proliferation rate^59–61^. Cell adhesion through integrins is essential for the G1/S cell cycle transition. Mechanistically, the activation of both integrins and growth factor receptors is required to get a sustained activation of Akt and Erk pathways, which in turn leads to cyclin D1 synthesis^62,63^. Hence, the regulation of integrin expression by the ECM proteins observed here is likely to also result in changes in cellular phenotypes. Follow-up mechanistic studies are required to further investigate the role of these integrin modulations in response to ECM proteins. For example, we can genetically modulate these integrins and observe the functional response of the cells to the corresponding ECM proteins.

MIA PaCa-2 responded to the ECM proteins differently when the cells were cultured in 2D and 3D systems **(Figure 5)**. There are multiple possible mechanisms behind these differential responses. For example, the 3D culture might promote cell aggregation that leads to an increase in cell-cell junctions. Similar to the integrins, the activation of cell-cell junction proteins, like E-cadherin, also regulates multiple pathways^64–66^. Moreover, the signaling from the integrin-ECM and the adherens junctions are known to compete with each other in dictating cellular phenotypes^67^. Hence, the increase in adherens junction in the 3D culture system might suppress the integrin-ECM signaling and diminish the cellular response towards ECM proteins. To test this, E-cadherin can be suppressed genetically or through TGF-β treatment, which will then shift the regulatory balance toward the integrin-ECM signaling. Further mechanistic and functional studies are required to fully elucidate the effects of ECM proteins on PDAC cells cultured in 3D.

Differences between paired primary and metastatic PDAC tumors have been reported at multiple levels, ranging from the genetic and epigenetic variations^68–71^ to the differences in cellular composition within the tumor microenvironment^72,73^, and even the mechanical properties of the tumor itself^74^. Our findings add another aspect of these differences, where the primary and metastatic PDAC cells may respond to the ECM proteins differently. However, it is important to note that the cell lines used in this study are derived from different patients, and there is a possibility that the differences observed here are driven by patient-to-patient variations. Nevertheless, our findings provide the premise for future studies to further investigate the differential responses to ECM proteins by using PDAC models with the same genetic backgrounds, such as the use of paired primary and metastatic tumor-derived PDAC organoids, genetically engineered mouse models of metastatic PDAC, or orthotopic mouse models with the same PDAC cell line. With these models, we can also investigate the driving mechanisms behind the genotype and phenotype differences between the primary and metastatic tumors. Is it driven by genomic instability that results in mutations that enable metastasis? Is it caused by other processes along the metastasis progression, such as confined cell migration that can lead to increased genomic instability^75^? Is it driven by the new microenvironment of the metastatic site, such as the presence of different soluble factors like growth factors and hormones, or even the changes in ECM composition and stiffness that have been shown to drive epigenetic changes^76^? Finally, considering that cell adhesion and spreading depend on ECM ligand density ^30,31^, and also the biphasic nature of the response ^32–35^, it is also important to note that the observations in this study are limited to the range of ECM concentrations that were used to coat the culture surface, and different responses might be observed when other ECM concentrations are used.

## Experimental procedures

### Cell culture

MIA PaCa-2 and PANC-1, the primary tumor-derived PDAC cell lines, were cultured in DMEM high-glucose media (Genesee Scientific). AsPC-1 and Capan-1, the metastatic PDAC cell lines derived from the ascites^36^ and the liver^37^, respectively, were cultured in RPMI 1640 media (Genesee Scientific). Both media were supplemented with 10% FBS (Sigma-Aldrich) and 1% penicillin and streptomycin (Corning). For experiments to study the effects of ECMs, 2% B27 (ThermoFisher) was used in place of FBS. Both the PDAC and glioblastoma organoids, HCM-CSHL-0092-C25 (ATCC PDM-39) and HCM-BROD-0103-C71 (ATCC PDM-121) respectively, are part of the Human Cancer Models Initiative and were acquired from ATCC. Maintenance of the organoid culture was performed following the instructions from ATCC, as described previously^77^.

### ECM proteins coating

Unless stated, collagen I (0.1mg/ml, Corning), fibronectin (10μg/ml, Sigma-Aldrich), laminin (10μg/ml, Corning), and vitronectin (2μg/ml, Sigma-Aldrich) were coated on plates according to manufacturer description. In short, Plates were incubated with collagen I overnight at 4°C; with fibronectin and laminin for 1 hour at room temperature; and with vitronectin at 37°C for 2 hours, after which it was transferred to 4°C overnight. After incubation, excess ECM proteins were removed, and the plates were air-dried at room temperature for 1 hour.

### Cell proliferation and attachment assay

For the proliferation assay, cells were seeded at 3200–6400 cells/cm^2^ and cultured for 6–12 days, and their numbers were measured every 2–4 days using the CellTitre-Glo 3D cell viability assay (Promega), as described by the manufacturer. Briefly, cells were seeded onto a white 96-well plate (Greiner Bio-One) coated with different ECM proteins, and cultured for 2, 3, 4, 6, 8, or 12 days. The CellTitre-Glo reagent was added to the wells 1:1. The plate reader (BioTek Synergy H1MF multi-mode) then shook the plate for 2 minutes with a delay time of 10 minutes, after which the luminescence was measured and recorded. The number of cells was extrapolated from the standard curve generated from the luminescence readings.

For the attachment assay, 8000 cells were seeded onto each well of the 96-well plate coated with different ECM proteins, and cultured for 6 or 16 hours for MIA PaCa-2 or Capan-1 cells, respectively. The wells were washed with PBS to remove the non-adherent cells, followed by fixation by 4% formaldehyde (EMS) for 10 minutes, DNA staining by 2μg/ml DAPI (Sigma-Aldrich) for 15 minutes, and fluorescence imaging to count the attached cells.

### Immunofluorescence and image processing

Cells/hydrogels were fixed in 4% formaldehyde, after which they were permeabilized in 0.5% Triton X-100 (MilliporeSigma) for 10 minutes and blocked with 5% bovine serum albumin (VWR) for 30 minutes. For nuclei and F-actin staining, cells were incubated in 2μg/ml DAPI and phalloidin-tetramethylrhodamine B isothiocyanate (Sigma-Aldrich) respectively, for 1 hour. Images were taken with an Olympus IX71 with an sCMOS camera (Prime 95B, Photometrics) and 10x/0.3NA, 20x/0.75NA, or 40x/0.6NA objective. All the image analysis was done with ImageJ^78^.

### Gemcitabine treatment

Cells were plated in either an ECM-coated or regular white-bottom 96-well plate and cultured for at least 24 hours prior to the gemcitabine treatment (MilliporeSigma). After 72 hours of gemcitabine treatment, the cell number was measured using the luminescence cell viability assay, as described above. To derive the gemcitabine dose-response curve, the luminescence data was fitted with the following equation^42^:

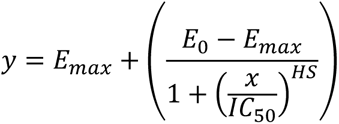

where *x* is the gemcitabine concentration, *y* is the corresponding luminescence readout, *E0* and *Emax* are the top and bottom asymptotes of the curve, respectively, *IC50* is the gemcitabine concentration at the half-maximal effect, and *HS* is the Hill slope coefficient. The Hill slope coefficient was constrained to -1 to assume that gemcitabine binding follows the law of mass action. Curve fitting and calculation of IC50 and Emax were done using GraphPad Prism v.10 (GraphPad).

### Transcriptomics sample preparation and analysis

After culturing the cells on different ECMs, RNA was isolated using the RNeasy Plus Micro Kit (QIAGEN). An RNA-seq library was prepared using the NEBNext Ultra II Directional RNA Library Prep Kit for Illumina (NEB) following the manufacturer’s instructions. The library was sequenced on the Novaseq 6000 at Florida State University Translational Science Laboratory, yielding ∼20 million reads per sample. The quality of the fastq reads from the sequencing was ascertained with FastQC (version 0.11.9) ^79^, after which they were aligned with the hg19 human reference genome using STAR (version 2.7.9a) ^80^. Library sizes were scaled and normalized with the median of ratios using the DESeq2 (version 1.32.0) ^44^. Differential expression was estimated using the result function in DESeq2. Adjusted p-values, using the Benjamini–Hochberg method, were computed and used to filter for significant genes. The heatmap showing the differential gene expression was made using the Pheatmap package in R ^81^. Specifically, significant differentially expressed genes (adjusted p-value < 0.05, abs(log2FC) > 0.58) in all of the ECM proteins were combined and clustered by K-mean (center = 4). Hierarchical clustering was used to cluster the column using Pearson’s correlation and Ward.D2. Pathway analysis was done using Enrichr ^45^. The significant pathways were filtered using FDR < 0.1.

For the transcriptomic-based gemcitabine sensitivity analysis, the list of genes that are positively and negatively correlated to gemcitabine sensitivity in PDAC patients was acquired from a previous study by the Tuveson Lab^46^. To visualize the gene expression of these genes, heatmap and hierarchical clustering of the normalized read counts were performed as described above. To calculate the sensitivity score, first, the normalized read counts of the negatively correlated genes were inverted by multiplying the values by -1, then the sensitivity score was calculated as an average of the normalized read counts from all correlated genes, both positively and negatively correlated genes.

### 3D PEG hydrogel culture

Functionalized PEG gel precursors (PEG-NDG-Lys, PEG-Gln) were made as described previously ^51^. Briefly, the PEG was functionalized by incubating the 10 kDa 8arm PEG vinyl sulfone (JemKem Technology) with Ac-FKGG-GDQGIAGF-ERCG-NH2 (TG-NDG-Lys) or H-NQEQVSPL-ERCGNH2 (TG-Gln) peptides (GenScript) in triethanolamine (0.3 M, pH 8.0) at 37°C for 20 minutes. The resulting PEG-NDG-Lys and PEG-Gln solutions were dialyzed using a 10,000 MWCO Slide-A-Lyzer dialysis cartridge (ThermoFisher) and lyophilized. The resulting lyophilized precursors were dissolved and reconstituted in deionized water to make a 13.33% w/v stock solution. To make a 10μl gel, 1μl of TBS (50 mM, pH 7.6); 1μl of collagen I (3.6mg/ml, Corning), fibronectin (1mg/ml, Sigma-Aldrich), laminin (1mg/ml, Corning), or vitronectin (0.5mg/ml, Sigma-Aldrich); and 1000 cells were added to 4μl of 10% PEG-NDG-Lys + PEG-Gln solution and 1μl distilled water. After the addition of 1μl of thrombin-activated FXIII (Corifact, CSL Behring), which triggers hydrogel formation, the well plate was flipped and incubated at 37°C for 30 minutes to allow for gelation.

### Statistical analysis

The figure legends specify the sample size for each condition. Statistical analyses were conducted using one-way analysis of variance (ANOVA) with post hoc Tukey-Kramer tests for the cellular morphology, attachment, proliferation, and gemcitabine sensitivity studies. The analyses were performed using GraphPad Prism v.10. All statistical tests were considered significant if *p* < 0.05. Unless otherwise stated, all plots show mean ± SEM.

## Acknowledgment

The authors in this study were supported by startup funds from Florida State University, awards from the Florida Department of Health’s Bankhead-Coley Cancer Research Program (award number 21B11) and Live Like Bella Pediatric Cancer Research Initiative (award number 23L06), and a collaborative Trans-Network Project linked to the Physical Science-Oncology Project 5U01 CA214282 from the National Cancer Institute of the National Institutes of Health. S. Ramakrishnan would like to acknowledge funding from the NSF FAMU CREST center award # 1735968 for this work. Corifact was a gift from CSL Behring. Pancreatic cancer and glioblastoma organoid lines are from the Human Cancer Models Initiative, https://ocg.cancer.gov/programs/HCMI. The authors would like to thank Dr. Terra Bradley for the careful editing of the manuscript.

---

**Figure S1.**
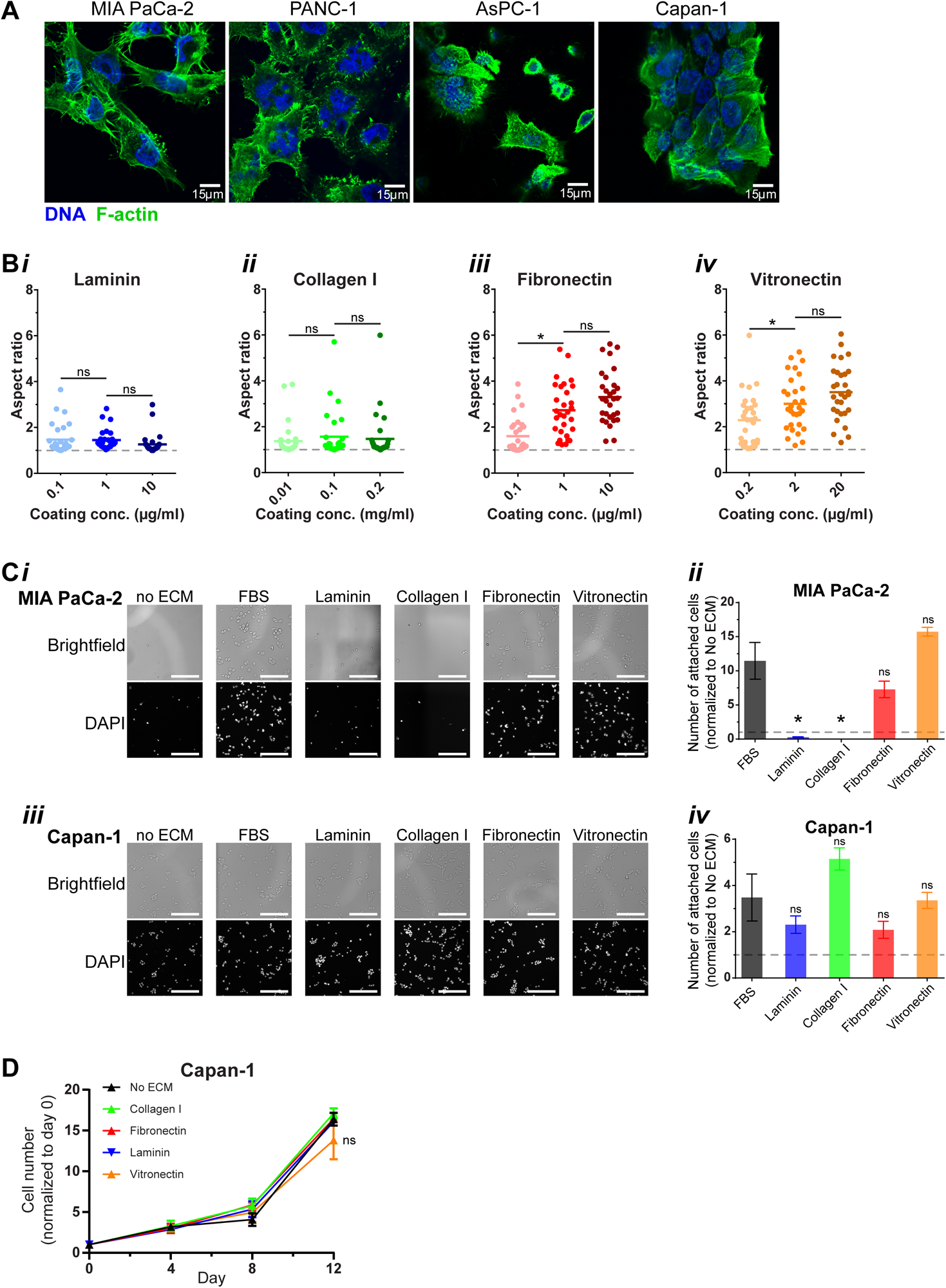
Both cellular morphology and attachment capability of MIA PaCa-2 cells are modulated by ECM proteins, but not for Capan-1 cells. **(A)** Images of MIA PaCa-2, PANC-1, AsPC-1, and Capan-1 stained with DAPI (blue) and Phalloidin (green) (scale bar: 15µm). **(B)** MIA PaCa-2 cells were cultured on surfaces coated with varying concentrations of laminin, collagen I, fibronectin, and vitronectin for 6 days. The degree of cell spreading was quantified by measuring the aspect ratio of the cells. For both laminin and collagen I coated surface, the cellular aspect ratio is close to 1 in all coating concentrations (**i-ii**), indicative of rounded cellular morphology, as shown in Fig. 1A. For both fibronectin and vitronectin-coated surfaces, a significant increase of aspect ratio is observed between the lower and the middle coating concentrations, however, the differences between the middle and the high concentrations are not statistically significant (**iii-iv**) (N = 2 experiments, n = 20-30 cells per group, \**p* < 0.05, ns: *p* > 0.05). **(C)** MIA PaCa-2 and Capan-1 cells were seeded onto the different surfaces and cultured for 6 and 16 hours, respectively, to allow for cell attachment. The cultures were washed with PBS to remove the non-adherent cells, followed by fixation, DNA staining, and counting of the attached cells. Representative images showing the cellular density of MIA PaCa-2 (**i**) cultured on the different ECM coatings. The cell counts were normalized to the number of attached cells in the no ECM group. For the MIA PaCa-2 cells, the number of attached cells in the laminin and collagen I groups is significantly lower when compared to the cells cultured in FBS media. While the cell count of the fibronectin and vitronectin groups are comparable to the FBS group (**ii**). For the Capan-I cells, the number of attached cells is statistically similar between the FBS group and all of the ECM coating groups (**iii-iv**) (N = 3 experiments, n = 3 wells per group, \**p* < 0.05 vs. FBS, ns: *p* > 0.05 vs. FBS, scale bar: 250µm). **(D)** The number of Capan-1 cells cultured on the different ECM coatings was quantified on days 4, 8, and 12, and normalized to the number of cells seeded. The proliferation rate of the cells cultured on the different ECM proteins was not significantly different from those cultured on no ECM plates (mean ± SEM, ns: *p* > 0.05 vs. no ECM).

**Figure S2.**
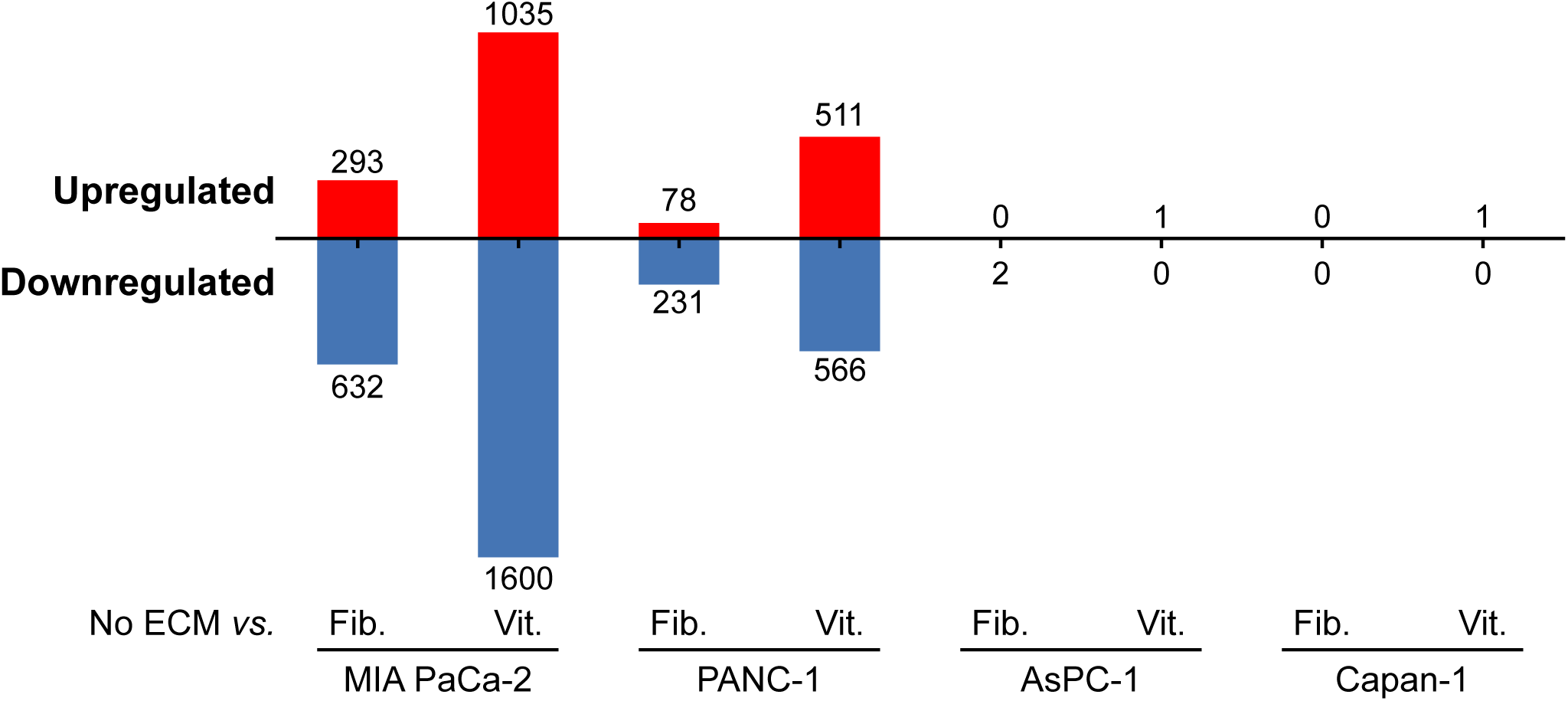
The fibronectin and vitronectin-driven transcriptomic modulation was observed in the primary tumor-derived cell lines, but not in the metastatic cell lines. MIA PaCa-1, PANC-1, AsPC-1, and Capan-1 cells were cultured on surfaces without ECM coating (no ECM) and surfaces with fibronectin and vitronectin coatings, then subjected to RNA-seq analysis. The bar plot shows the number of differentially expressed genes (vs. no ECM) for each cell line. A high number of regulated genes (78-1600 genes) were observed in both of the primary tumor-derived cell lines, MIA PaCa-2 and PANC-1, while a limited number (0-2 genes) were observed in the metastatic lesion-derived lines, AsPC-1 and Capan-1.

**Figure S3.**
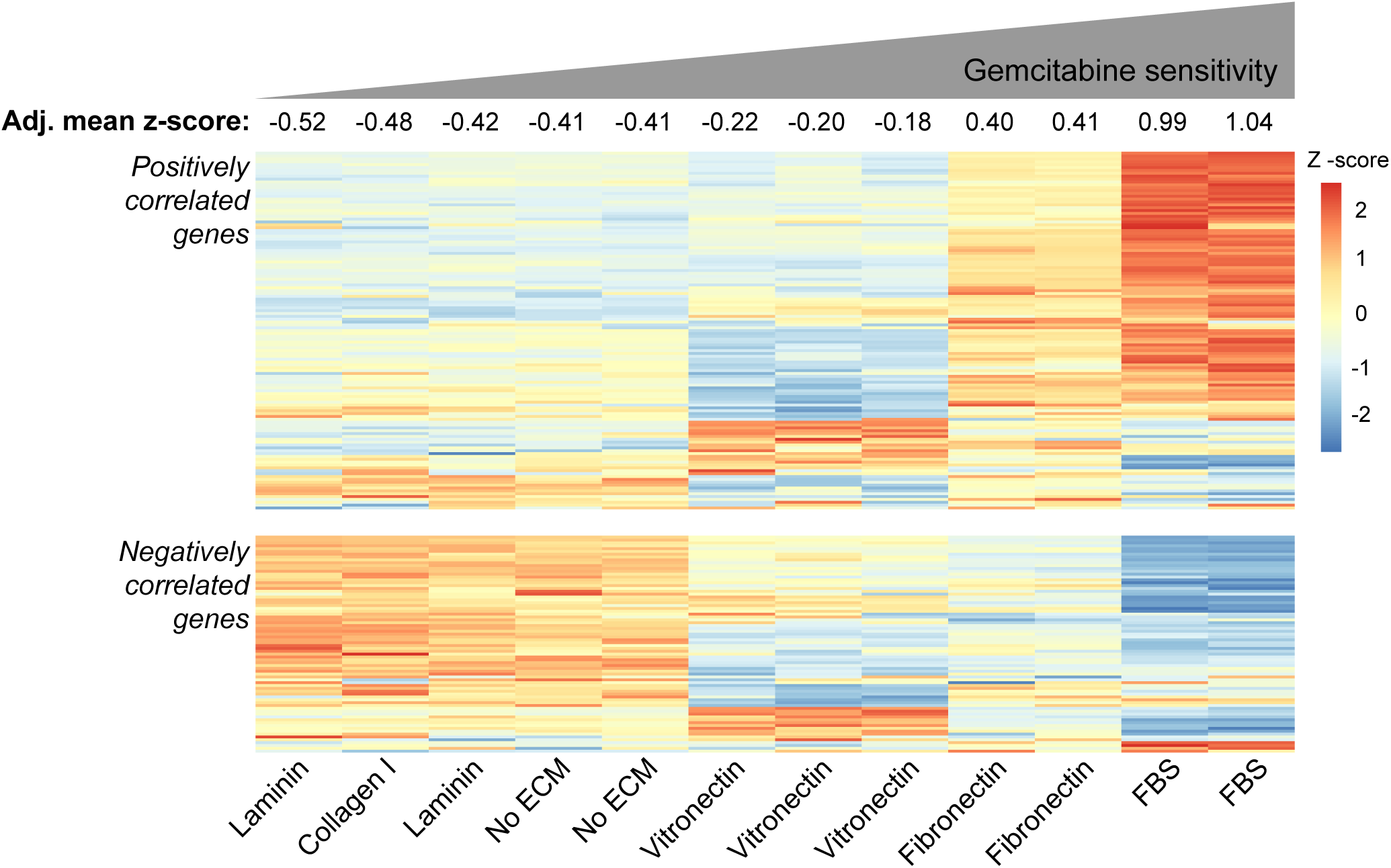
The transcriptomic-based gemcitabine sensitivity predictions confirm the modulation of gemcitabine sensitivity by the ECM proteins in MIA PaCa-2 cells. Heatmap showing normalized read counts of the genes that are positively (top) and negatively (bottom) correlated with gemcitabine sensitivity in PDAC patients^46^. The genes that positively correlated with gemcitabine sensitivity are generally upregulated in the FBS and fibronectin groups, while the negatively correlated genes are upregulated in the no ECM, laminin, and collagen I groups. To get the sensitivity score (adjusted mean z-score), the normalized read counts of the negatively correlated genes were inverted, then the sensitivity score was calculated as the average of normalized read counts from all correlated genes. High positive scores indicate higher sensitivity to gemcitabine treatment. Based on the sensitivity score, the analysis predicts that the FBS group is the most sensitive to gemcitabine treatment, followed by fibronectin, vitronectin, and the other groups with rounded morphology: no ECM, laminin, and collagen I. This prediction confirms the findings in Fig. 2C.

**Figure S4.**
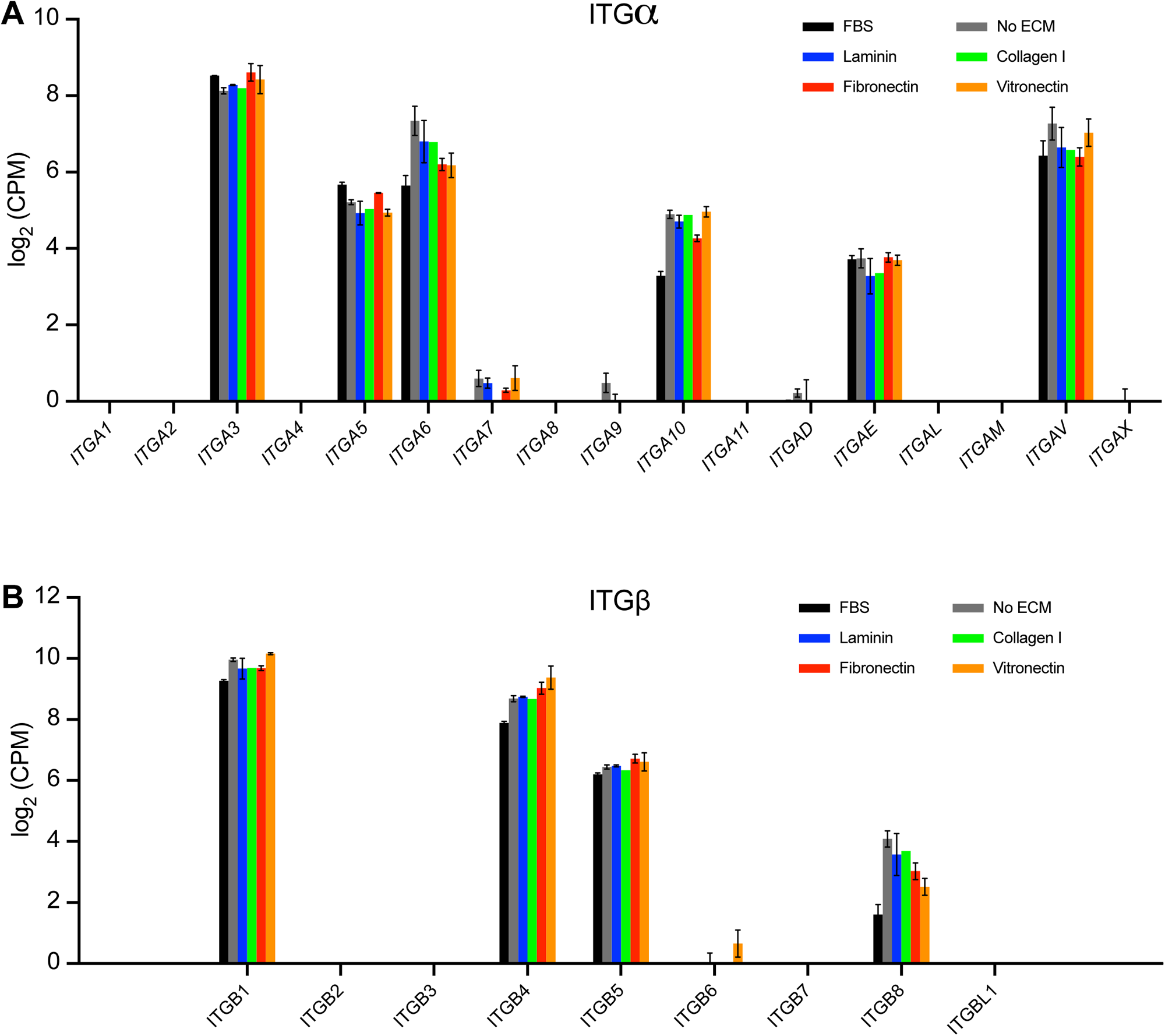
Integrin gene expressions on different ECM proteins. **(A–B)** Bar plots showing the normalized read counts of ITGα and ITGβ in MIA PaCa-2 cells cultured on different ECM proteins.

**Figure S5.**
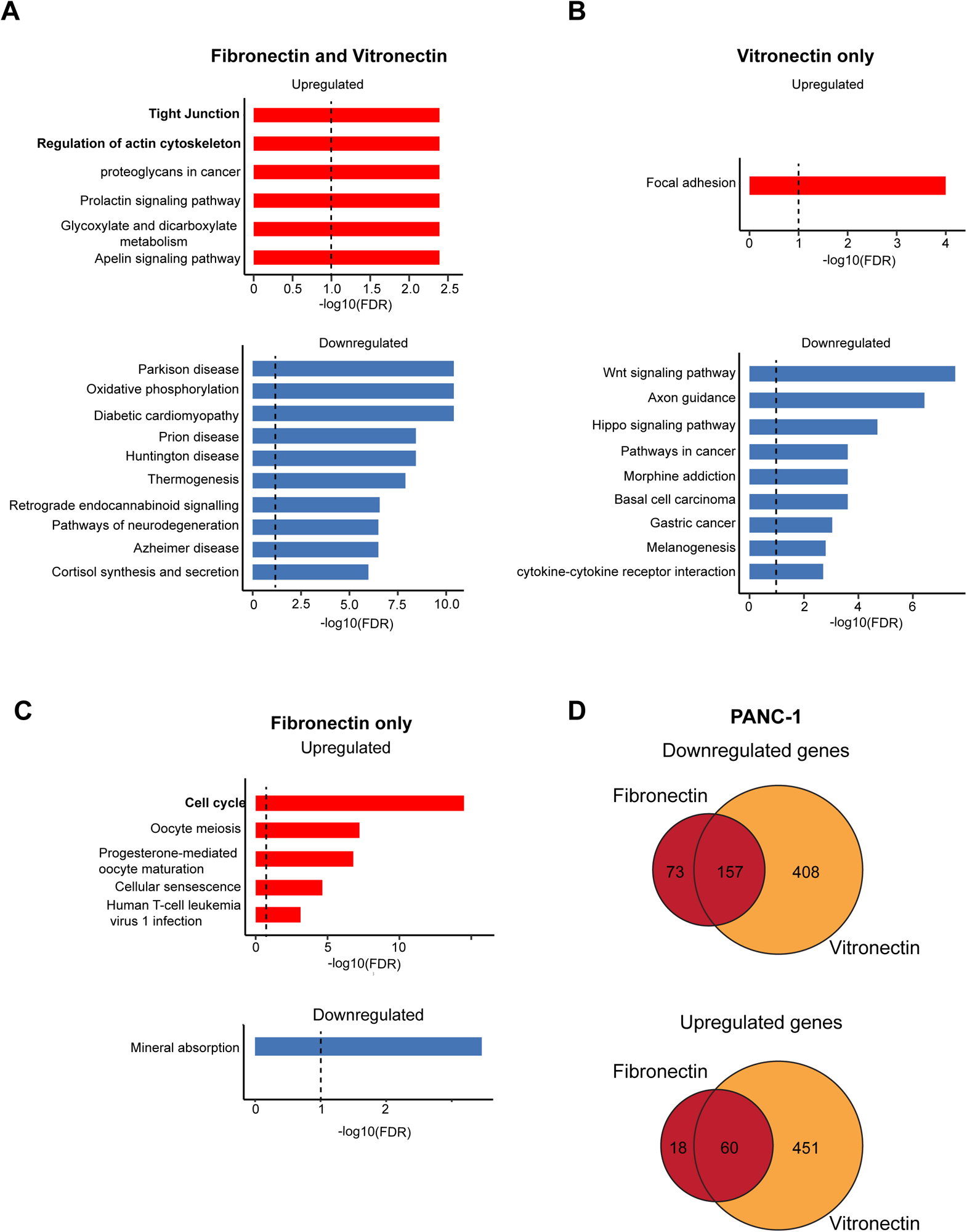
Differential regulation of MIA PaCa-2 and PANC-1 when cultured on fibronectin and vitronectin. **(A–C)** Bar plots showing the KEGG pathways that are enriched by the genes regulated in both fibronectin and vitronectin, vitronectin, and fibronectin only. The genes in the overlap of the Venn diagrams in Fig. 4A were subjected to pathway analysis using Enrichr and a maximum of the top 10 significant KEGG pathways were selected (FDR < 0.1). The red and blue bars indicate upregulated and downregulated pathways respectively. **(D)** From the PANC-1 transcriptomic data, the Venn diagrams show the number of differentially expressed genes (vs. no ECM) that are regulated in both fibronectin and vitronectin (overlap), and also the genes that are unique to fibronectin or vitronectin. The left panel **(i)** shows that 157 genes are upregulated in cells cultured on fibronectin and vitronectin, while 408 and 73 genes are uniquely upregulated in vitronectin and fibronectin respectively. The right panel **(ii)** shows that 60 genes are downregulated in cells grown on fibronectin and vitronectin, while 451 and 18 genes are downregulated uniquely in vitronectin and fibronectin respectively.

